# STIM1 signals through NFAT independently of Orai1 and SOCE to regulate breast cancer cell migration

**DOI:** 10.1101/2022.10.23.513385

**Authors:** Ayat S. Hammad, Fang Yu, F. David Horgen, Khaled Machaca

**Affiliations:** College of Health and Life Science, Hamad bin Khalifa University, Doha, Qatar; Calcium Signaling Group, Research Department, Weill Cornell Medicine Qatar, Doha, Qatar; Department of Physiology and Biophysics, Weill Cornell Medicine, New York, NY, USA; Department of Natural Sciences, Hawaii Pacific University, Honolulu, Hawaii, USA

## Abstract

Store-operated calcium entry (SOCE) contributes to several physiological and pathological conditions including transcription, secretion, immunodeficiencies, and cancer. SOCE has been shown to be important for breast cancer cell migration where knockdown of SOCE components (STIM1 or Orai1) decreases cancer metastasis. Here we show unexpectedly that STIM1 knockout (KO) metastatic MDA-MB-231 breast cancer cells migrate faster and have enhance invasion capacity compared to parental cells. In contrast, Orai1-KO cells, which have similar levels of SOCE inhibition as STIM1-KO, migrate slower than the parental cell line. This shows that the enhanced migration phenotype of STIM1-KO cells is not due to the loss of a Ca^2+^ entry through SOCE, rather it involves transcriptional remodeling. Interestingly, NFATC2 is significantly downregulated in STIM1-KO cells and overexpression of NFATC2 reversed the enhanced migration of STIM1-KO cells. This demonstrates that STIM1 modulates NFATC2 expression independently of its role in SOCE.

**SUMMARY STATEMENT:** Breast cancer cells migrate faster when the ER Ca^2+^ sensor STIM1 in knocked out due to downregulation of NFAT1 expression independent of Ca^2+^ influx.

## INTRODUCTION

Breast tumors are the most prevalent type of cancer among females accounting for over 27% of all cancer incidence in women (2020). Over 90% of all cancer deaths are due to metastasis rather than the primary tumor (Hanahan and Weinberg, 2000). Cell migration is important for cancer cell metastasis and depends on Ca^2+^ signaling in particular store-operated Ca^2+^ entry (SOCE) (Hammad and Machaca, 2021).

SOCE is mediated by interactions between the ER Ca^2+^ sensor STIM1 with the Orai channel family at the PM (Prakriya and Lewis, 2015). Store depletion following agonist-dependent Ca^2+^ release from ER Ca^2+^ stores results in clustering of STIM1 at ER-PM junctions, where it recruits and gates Orai1 allowing Ca^2+^ influx. SOCE has been shown to be critical for breast cancer metastasis as a reduction of either STIM1 or Orai1 in the metastatic MDA-MB-231 cells decreases tumor metastasis (Yang et al., 2009). Supporting these findings, overexpression of STIM1 in the less aggressive MCF-7 breast tumor cells boosts their migration capacity (Zhang et al., 2017). Indeed, for the majority of cell lines tested, both cancerous (lung, breast, prostate, colorectal, etc.) or not (HEK293, VSMC, MEF), downregulation of SOCE by lowering STIM1 or Orai1 levels or function decreases cell migration capacity (recently reviewed (Hammad and Machaca, 2021)). There are however a few exceptions. Complete knockout of STIM1 in MEFs or overexpression of a dominant negative STIM1 in osteosarcoma cells enhances cell migration (Huang et al., 2018; Lin et al., 2021). Furthermore, in the context of cancer metastasis, recent evidence suggests that SOCE suppression is important in melanoma metastasis (Gross et al., 2022), arguing that the role of SOCE in cancer metastasis is intricate and depends on the cancer type.

The differential effect of SOCE in different cancer cell types could be due to the SOCE-dependent modulation of focal adhesions (FA). SOCE affects FA dynamics in a complex fashion in moving cells by strengthening nascent FA on the front and supporting disassembly of FA at the rear end to allow for directional cell movement (D’Souza et al., 2020; Shellard and Mayor, 2020; Tsai et al., 2014).

To better define the mechanisms involved in SOCE-dependent regulation of cell migration in metastatic breast cancer cells, we used CRISPR-dependent gene editing to engineer MDA-MB-231 breast cancer cells with specific SOCE gene knockouts, including STIM1, STIM2, STIM1 and 2, and Orai1. We show that the STIM1-KO cells migrate and invade faster independent of SOCE modulation but rather due to NFAT-dependent transcriptional remodeling.

## RESULTS AND DICUSSION

### Generation of STIMs and Orail KO MDA-MB-231 cell lines

As expected from the literature blocking SOCE with the specific inhibitor BTP2 reduces collective cell migration in the wound healing assay in two highly metastatic breast cancer cell lines MDA-MB-231 and MDA-MB-463 (Supplemental Fig. S1A-B). In contrast BTP2 had a less pronounced, if any, effect in the poorly metastatic MCF7 breast cancer cells (Supp. Fig. S1C), which expresses significantly lower levels of STIM1 compared to MDA-MB-231 (Kulkarni et al., 2019). This argues that SOCE plays a more important role in highly migratory metastatic breast cancer cell lines. To better define the mechanisms involved we deleted SOCE proteins using CRISPR/Cas9 genome editing in MDA-MB-231 cells. We engineered STIM1-KO (S1); STIM2-KO (S2); STIM1/2-KO (S1/S2); and an Orai1-KO (O1) lines and genotyped them by sequencing either genomic or mRNA (Table 1). Western blots confirm the specific loss of the protein of interest (Supp. Fig. S1D-F). Due to the lack of specific Orai1antibodies, we could not assess Orai1 protein levels, so we tested for loss of Orai1 functionally as discussed below. The gene knockouts did not affect cell proliferation in any of the KO cell lines using two different assays MTT and Alamarblue (Supp. Fig. 1G-H). Furthermore, knockout of STIM1, STIM2, or Orai1 did not alter Ca^2+^ store content (Supp. Fig. 1I). Resting Ca^2+^ levels were similar in all cell lines except for the STIM2-KO where it was slightly lower (Supp. Fig. 1J) as previously reported (Brandman et al., 2007). The effects of the knockouts on SOCE were as expected: compared to the parental MDA-MB-231 line the STIM1-KO, STIM1/2-KO, and Orai1-KO lines showed close to complete loss of SOCE in the Ca^2+^ re-addition assay following Ca^2+^ store depletion with thapsigargin (Fig. 1A-B), whereas the levels of SOCE in the STIM2-KO line were similar to those in control cells (Fig. 1A-B). We further validated the loss of SOCE in the STIM1-KO and Orai1-KO lines using agonist dependent Ca^2+^ signaling with ATP in Ca^2+^ containing medium. This resulted in an initial Ca^2+^ release followed by low level Ca^2+^ influx for extended periods of time in control cells (Fig. 1C). In the STIM1-KO and Orai1-KO lines the release phase was unaffected, but the extended influx phase was lost (Fig. 1C). Peak Ca^2+^ release levels were similar (Fig. 1C) consistent with no alterations in Ca^2+^ store content (Supp. Fig. 1I).

**Table 1.** Sequence validation of the knockout MDA-MB-231 cell lines

| Gene | Clone | Exon | HGVS and Affected Protein |
| --- | --- | --- | --- |
| **STIM1** | Clone 1 & 2 | 3 | NM_001277961:g.898delT; p.(Glu111Argfs*31) |
| **STIM2** | Clone 1<br>Clone 2 | 2 | NM_001169117:g.643_644delGT; p.(Val89Argfs*4)<br>g.632_639delGTGGAATT; p.(Gly86Serfs*5) |
| **STIM1/2** (from<br>STIM1-KO) | Clone 1<br>Clone 2 | 2 | NM_001169117:g.640_641delGA; p.(Glu88Serfs*5)<br>g.642_646delAGTAG; p.(Val89Glyfs*3) |
| **Orai1** | Clone 1<br>(compound heterozygous) | 2 | NM_032790:g.633_634delAC; p.(Ile148Argfs*18)<br>g.636_641delTCGAGG; p.(Ile148_Ala150delinsThr) |

**Figure 1.**
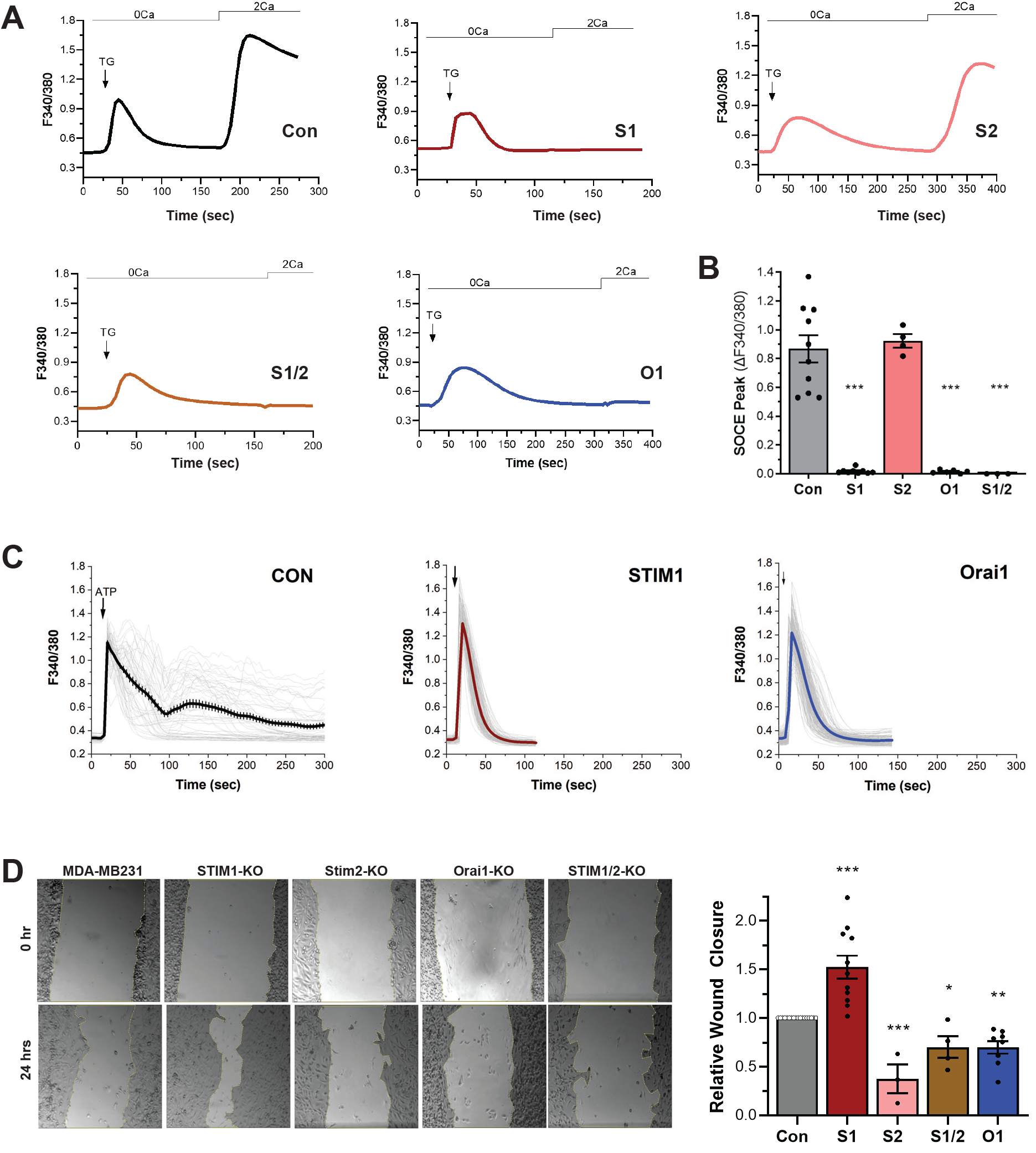
STIM1 knckout inhibits SOCE and enhances sheet cell migration in MDA-MB-231 cells. **(A)** SOCE example traces from the parental MDA-MB-231 cell line (Con) and the different knockout cell lines: STIM1 (S1), STIM2 (S2), STIM1 and STIM2 (S1/2), and Orai1 (O1). Ca^2+^ stores were depleted with thapsigargin (TG, 1 μM) in Ca^2+^-free media followed by addition of 2 mM ca^2+^ as indicated. **(B)** Summary of SOcE peak values in the different cell lines as indicated (Mean ± SEM, n=3-10, ANOVA with Tukey correction, *** p<0.001). **(C)** Cells were treated with ATP (20 μM) in Ca^2+^ containing media. Individual cell responses are shown in light gray with the Mean ± SEM as dark traces (n=51-181 cells). **(D)** Example images of wound closure over a 24 hrs time period for the different cells as indicated (left panel) and summary of relative wound closure compared to the parental MDA-MB-231 cells (Con) (right panel, Mean ± SEM, n=3-18, ANOVA with Fisher LSD, *** p<0.001, ** p < 0.01, * p < 0.05).

### STIM1 knockout enhances cell migration

We then tested cell migration and invasion capacity of the different KO lines using the wound healing assay to evaluate collective cell migration, time lapse recordings to evaluate individual cell migration, and the Boydon Chamber assay with Matrigel as an ECM substitute to test for invasion. Consistent with an important role for SOCE in cell migration and cancer metastasis, Orai1-KO reduces collective cell migration (Fig. 1D). STIM2-KO also significantly reduces wound closure capacity (Fig. 1D). In contrast, STIM1-KO cells exhibited enhance collective cell migration and this was reversed in the STIM1/2-KO cells (Fig. 1D). This is surprising especially that STIM1-KO and Orai1-KO cells reduce SOCE to similar levels (Fig. 1A-C), arguing that the STIM1-KO effect is independent from SOCE inhibition.

Based on these results we wanted to further assess the requirement for other Ca^2+^ signaling pathways in the enhanced migration observed in STIM1-KO cells. Ca^2+^ signaling is polarized in migrating cells with a gradient across the cell (low in the front) due, at least in part, to enrichment of the plasma membrane Ca^2+^-ATPase (PMCA) at the front, which lowers Ca^2+^ levels (Tsai et al., 2014). Furthermore, migrating cells exhibit localized Ca^2+^ pulses at the front that depend on the stretch activated Ca^2+^ permeable TRPM7 channel (Visser et al., 2013; Wei et al., 2009). In addition, TRPM7 modulates cellular tension and focal adhesions in migrating cells (Middelbeek et al., 2012). To broadly test for the involvement of Ca^2+^ influx or extrusion pathways at the plasma membrane, we performed the wound healing assay on MDA-MB-231 cells treated BTP2 to specifically block SOCE, or high La^3+^ (1 mM) to non-specifically block Ca^2+^ influx and extrusion. BTP2 reduced cell migration by 38.3+24.5% whereas La^3+^ had a more dramatic effect with 76.6±0.19% reduction (Fig. 2A). In contrast, BTP2 did not inhibit cell migration in STIM1-KO cells and La^3+^ resulted in a smaller reduction of 45.8+20% (Fig. 2B). These data argue that the enhanced cell migration observed in STIM1-KO cells remains partially dependent on Ca^2+^ but less so then in the control MDA-MB-231 cells.

**Figure 2.**
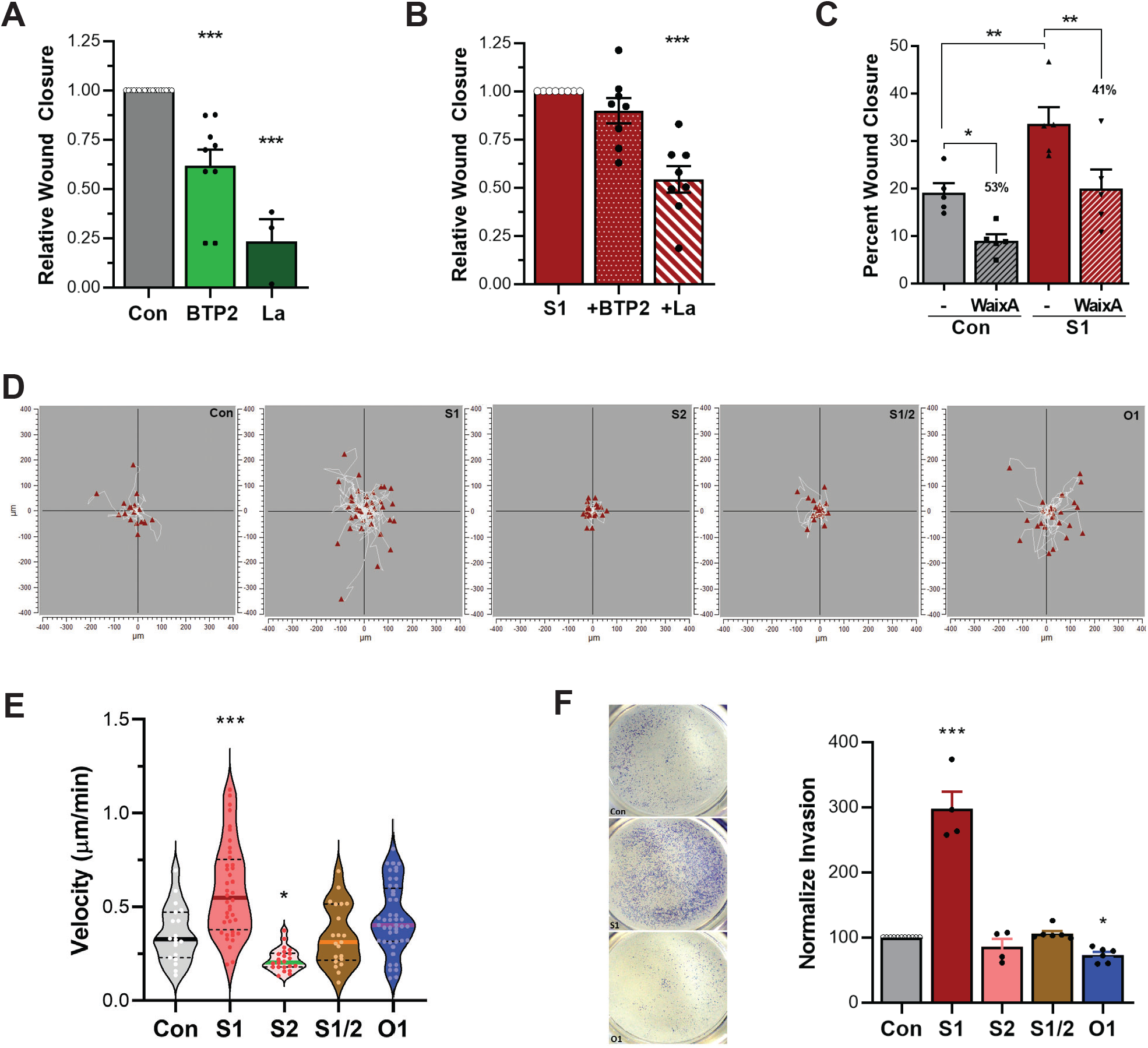
STIM1-KO enhances cell migration and invasion. **(A-B)** Inhibition of wound closure with BTP2 (2 μM) and LaCl_3_ (1 mM) at 24 hrs in MDA-MB-231 **(A)** and the STIM1-KO **(B)** cell lines. Mean ± SEM, n=3-18 for (A) and n=8 for (B), ANOVA with Dunnett correction, *** p<0.001. **(C)** Effect of the TRPM7 inhibitor waixenicin A (WaixA, 20 μM) on wound closure in MDA-MB-231 (Con) and STIM1-KO (S1) cells with the percent inhibition shown above the WaixA treatment (Mean ± SEM, n=5, ANOVA with Fisher LSD, ** p < 0.01, * p < 0.05). **(D-E)** Individual cells trajectories for the different genotypes as indicated **(D)** and the corresponding velocity **(E)**. Mean ± SEM, n=19-45, ANOVA with Fisher LSD, *** p<0.001, * p < 0.05. **(F)** Boyden chamber invasion assay for the different genotypes (Mean ± SEM, n=4-11, ANOVA with Fisher LSD, *** p<0.001, * p < 0.05).

We next tested whether TRPM7 supports the enhanced migration in STIM1-KO cells. We inhibited TRPM7 using the specific blocker waixenicin A (Visser et al., 2013). Waixenicin A reduced collective cell migration in both the control and STIM1-KO cell lines to similar extents, arguing against differential modulation of TRPM7 in the STIM1-KO cells (Fig. 2C).

Collective cell migration requires adjacent cells to maintain their cell-cell contact, which depends on coordinated signaling and regulation of actin dynamics (Friedl and Gilmour, 2009). To test whether the enhanced migration capacity of the STIM1-KO observed in the wound healing assay translates to individual cell migration, we quantified cell migration in time lapse video recordings for several hours. The trajectories of individual cells from the different genotypes is shown in Figure 2D. As is apparent from quantifying cell velocity, STIM1-KO cells migrate faster than the other cells, whereas STIM2-KO cells migrate slower (Fig. 2E). This shows that similar to collective cell migration, STIM1-KO leads to an enhanced individual cell migration.

During the process of metastasis, cancerous cells need to degrade the extracellular matrix (ECM) to reach the lymphatic or blood vessels through a process termed invasion. We thus tested the invasiveness in the Boyden chamber assay. Similar to individual and collective cell migration STIM1-KO cells exhibit enhanced invasion capacity compared to the parental MDA-MB-231 and other KO cell lines (Fig. 2F). Therefore, in three separate assays for cell migration and invasiveness STIM1-KO MDA-MB-231 cells consistently show enhanced migration and invasion capacity.

### Transcriptomic remodeling in STIM1-KO favors migration

Focal adhesions (FA) play a critical role in cell migration and are modulated by SOCE (Hammad and Machaca, 2021). Compared to the parental MDA-MB-231 cells, STIM1-KO cells show increased FA density and size in TIRF imaging (Fig. 3A-B). An increase in FA size was also observed in the STIM1/2-KO and decreased density in Orai1-KO cells (Fig. 3A-B). The increase in focal adhesion size has been previously documented in MDA-MB-231 cells following STIM1 or Orai1 knockdown (Yang et al., 2009). We observe a similar increase in FA size in the STIM1-KO cells but not in the Orai1-KO cells (Fig. 3C). Although these changes in FA density and size likely contribute to regulating cell migration as has been previously argued (Yang et al., 2009), they do not correlate with the observed migration phenotypes of the different KO cell lines and thus cannot account for the STIM1-KO enhanced migration.

**Figure 3.**
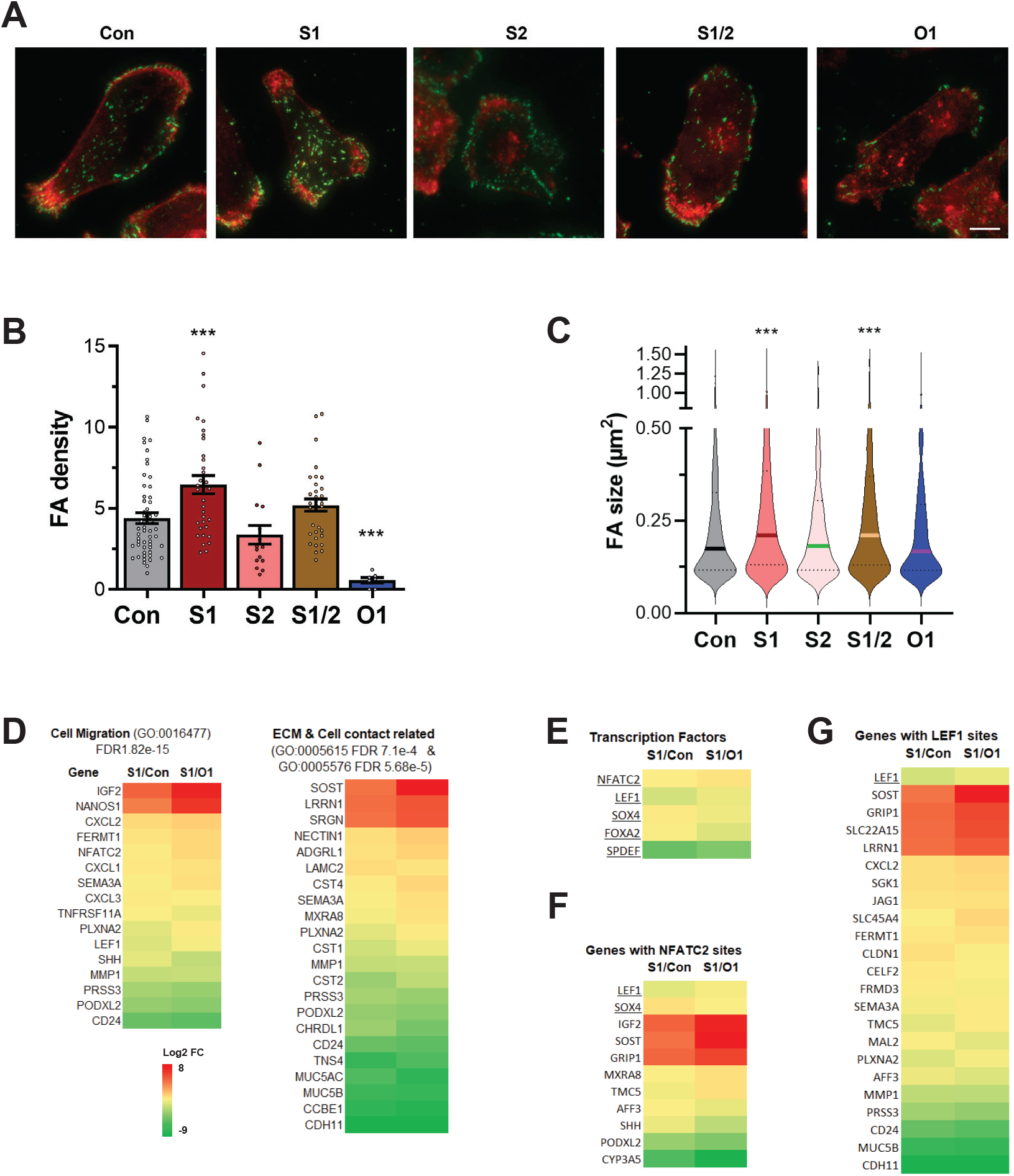
Focal adhesions and transcriptional remodeling in STIM1-KO cells. **(A)** Example TIRF images from cells stained for p-paxillin (green) and phalloidin to visualize actin (red). Scale bar is 10 μm. **(B)** Focal adhesion (FA) density for the different genotypes reported as the percent of the cell footprint occupied by focal adhesion (Mean ± SEM, n=7-55, ANOVA with Fisher LSD, *** p<0.001). **(C)** FA size (Mean ± SEM, n=467-1700, ANOVA with Fisher LSD, *** p<0.001, * p < 0.05). **(D)** Heat maps of Log2 fold changes of differentially expressed genes (DEG) between STIM1-KO and MDA-MB-MDA-MB-231 (S1/Con) or Orai1-KO (S1/O1) following pathway enrichment analysis with the enriched pathways indicated. **(E)** The transcription factors detected in the DEG set. **(F-G)** Genes that have an NFATc2 **(F)** or a LEF1 **(G)** consensus binding site in their promoter region.

We next performed transcriptomics analysis as an unbiased approach to identify differentially regulated genes in the STIM1-KO cells as compared to both the parental MDA-MB-231 (Con) and the Orai1-KO. We reasoned that if changes in gene expression modulate the enhanced migration of the STIM1-KO cells, these changes must be conserved in the STIM1-KO as compared to both the Con and Orai1-KO cells. We focused on genes that are highly up or down regulated with Log2 fold-change of ±2 and a p-adjusted (q value) significance of <0.05. Heat map and volcano plots of these differentially expressed genes (DEG) in the STIM1-KO compared to both Orai1-KO and Con cells are shown Supplemental Figure 2A-C. We confirmed the RNAseq data for a select number of genes by RT-PCR (Supp. Fig. 2D). Further pathway enrichment analysis of the DEG in STIM1-KO cells using String (string-db.org, v.11.5) identified genes involved in cell migration and related to the ECM and cell-ECM contact (Fig. 3D). This is consistent with the enhanced migration observed in STIM1-KO as compared to both the parental and Orai1-KO cell lines. The comparative analysis to Orai1-KO removes DEGs due to the loss of SOCE.

We then asked whether there are alterations in specific transcription factors that could explain the transcriptional remodeling observed in the STIM1-KO cells. Several transcription factors were significantly downregulated (Log2FC -2.5 to -6.5) in the DEG subset, including NFATC2 and LEF1 (Fig. 3E). NFATC2 and LEF1 were of particular interest because several of the genes that are specifically up or downregulated in STIM1-KO cells have either NFATC2 (Fig. 3F) or LEF1 (Fig. 3G) or both binding sites in their promoter and are thus candidates for expression regulation by these transcription factors. In fact, LEF1 is one of the candidate genes that could be regulated by NFATC2, suggesting the NFATC2 may be playing an important role in the remodeling of the transcriptome in the STIM1-KO cells.

We extended and validated the RNAseq data by assessing the expression levels of NFATC2 by Western blotting (Fig. 4A-B) and immunofluorescence (Fig. 4C-D). In both cases we observe a dramatic downregulation of NFATC2 protein levels in the STIM1-KO cells but not in the STIM2-KO or Orai1-KO cells (Fig. 4A-D). We further assessed the ability of NFATC2 to translocate to the nucleus in response to SOCE activation. As expected, given the normal levels of SOCE in the Con and STIM2-KO cells, NFATC2 translocates to the nucleus but not in the cell lines where SOCE was abrogated, STIM1-KO and Orai1-KO cells (Fig. 4E).

**Figure 4.**
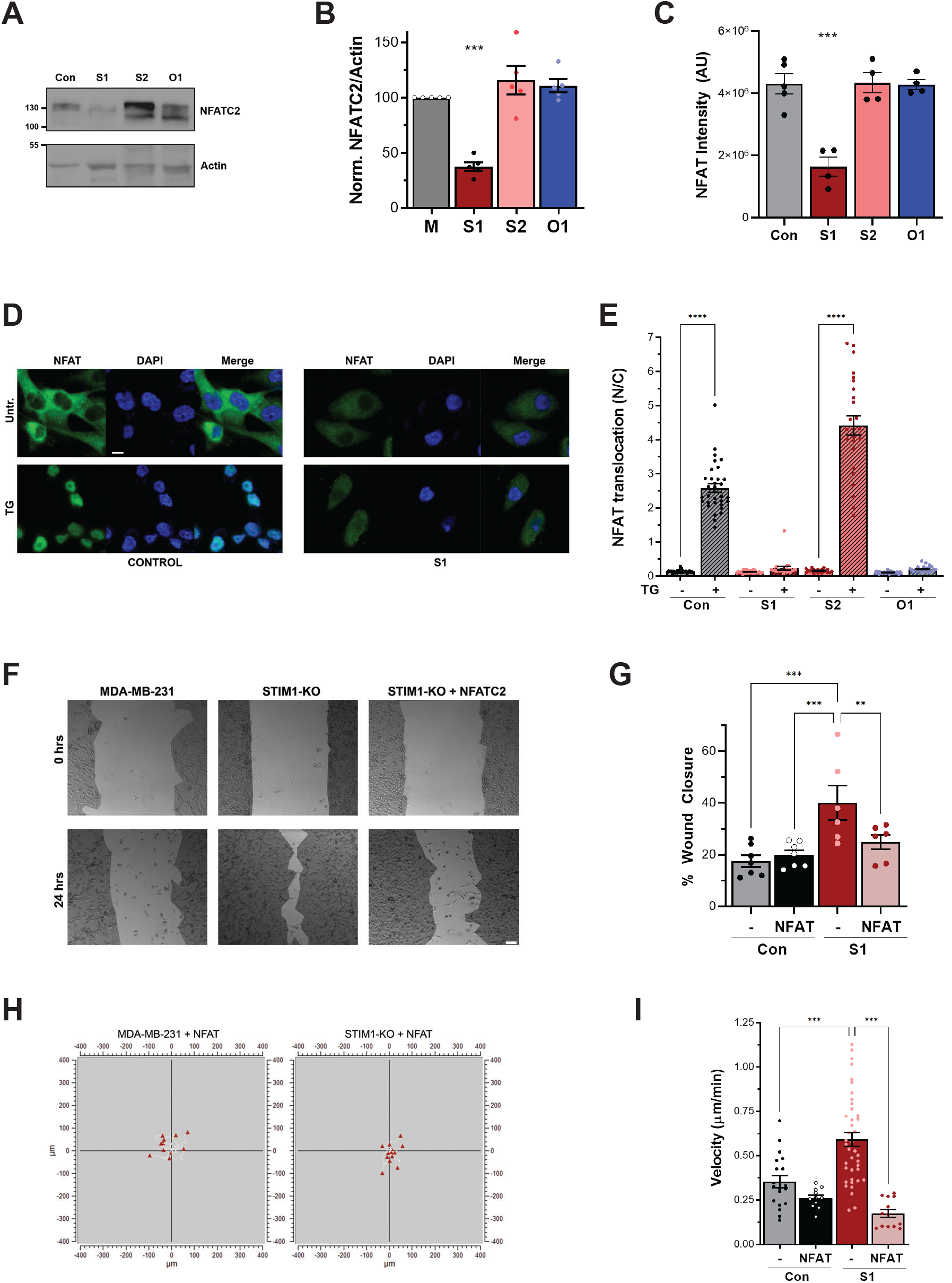
NFATC2 expression reverses the enhanced migration in STIM1-KO cells. **(A)** Western Blot of NFATC2 with actin as a loading control for the different cell lines as indicated. **(B)** Relative NFATC2 expression levels quantified from Western blots (Mean ± SEM, n=4-5, ANOVA with Fisher LSD, *** p<0.001). **(C)** Quantification of NFATC2 levels from immunostaining experiments as in (D). (Mean ± SEM, n=4-5, ANOVA with Fisher LSD, *** p<0.001). **(D)** Representative immunostaining experiment for NFATC2 before (Unt.) or after exposure to 1 μM thapsigargin in MDA-MB-231 (Control) and STIM1-KO cells (S1). Cells were also treated with Hoechst 33342 to stain nuclei. Scale bar is 10 μm. **(E)** Quantification of NFATC2 nuclear translocation measured as the ratio of NFATC2 localizing to the nucleus (N) over the cytoplasm (C) (Mean ± SEM, n=21-33, ANOVA with Tukey correction, *** p<0.001). **(F)** Example images from the wound healing assay from MDA-MB-231, STIM1-KO, and STIM1-KO overexpressing NFATC2. Scale bar is 100μm. **(G)** Quantification of wound closure (Mean ± SEM, n=6-7, ANOVA with Fisher LSD, *** p<0.001, ** p<0.01). **(H)** Individual cells trajectories for MDA-MB-231 and STIM1-KO cells overexpression NFATC2, and **(I)** the corresponding velocity (Mean ± SEM, n=13-40, ANOVA with Fisher LSD, *** p<0.001).

### NFATC2 overexpression rescues the enhanced migration in STIM1-KO cells

If the faster migration observed in the STIM1-KO cells is due to transcriptional remodeling due to, at least in part, NFATC2 downregulation, then overexpression of NFATC2 should reverse this phenotype. We used lentiviral infection to express NFATC2 in control and STIM1-KO cells and tested collective cell migration in the wound healing assay. NFATC2 expression reversed the rapid migration phenotype observed in STIM1-KO to levels similar to those observed in the parental cell line (Fig. 4F-G). Importantly, expression of NFATC2 in MDA-MB-231 cells did not slow down their collective cell migration (Fig. 4G), showing that the reversal of enhanced STIM1-KO cell migration is not due to a non-specific effect due to over-expression of NFATC2. We further assessed individual cell migration following NFATC2 expression, and similar to the wound healing assay the enhanced cell migration velocity in STIM1-KO cells was reversed in STIM1-KO cells expressing NFATC2 (Fig. 4H-I), without a reduction in cell velocity in control cells expressing NFATC2 (Fig. 4I). Collectively these data show that NFATC2 expression is sufficient to reverse the enhanced migration of STIM1-KO cells.

The canonical function of STIM1 is a resident ER Ca^2+^ sensor that partners with the Orai1 Ca^2+^ channel to activate SOCE. However, additional SOCE-independent roles for STIM1 are emerging in the literature, including the regulation of ion channels other than the Orai1 family (Mignen et al., 2007; Park et al., 2010; Wang et al., 2010; Zeng et al., 2008); modulation of the activity of plasma membrane and ER Ca^2+^ ATPases (Lee et al., 2014; Ritchie et al., 2012); regulation of adenylyl cyclases that control cAMP levels (Martin et al., 2009; Motiani et al., 2018); and maintenance of ER-PM contact sites in contractile VSM cells (Krishnan et al., 2022). Here we unravel a novel function of STIM1 as an inhibitor of NFATC2 transcription in metastatic breast cancer cells. When STIM1 is completely knocked out this leads to a dramatic downregulation of NFAT expression resulting in transcriptional remodeling and the associated enhanced migration.

Consistent with previous findings pharmacological inhibition of SOCE or Orai1-KO reduces cell migration using multiple assays in the metastatic MDA-MB-231 cells. In contrast, STIM1-KO MDA-MB-231 cells migrate significantly faster. The enhanced migration of STIM1 cells is independent of SOCE inhibition or Orai1 and is rather due to transcriptional remodeling, which affects genes involved in cell adhesion and migration. Interestingly, the transcription factor NFATC2 (NFAT1) is significantly downregulated in STIM1-KO cells. This is surprising as NFAT expression was not altered in Orai1-KO cells, which exhibited loss of SOCE to similar levels to those observed in STIM1-KO cells. NFATC2 downregulation is functionally significant as expressing NFATC2 was sufficient to reverse the enhanced migration phenotype observed in STIM1-KO cells, without any effect in control cells. Collectively our data reveal a functional link between STIM1 and NFATC2 expression that is independent of Ca^2+^ influx through SOCE but requires the STIM1 protein.

These findings are surprising because NFAT is one of the canonical effectors downstream of SOCE and specifically activates in response to Ca^2+^ influx through SOCE over other Ca^2+^ influx and release pathways (Kar et al., 2021; Oh-hora and Rao, 2008). Ca^2+^ influx through SOCE activates the dual specificity phosphatase calcineurin in the SOCE microdomain, which is turn dephosphorylates and activates NFAT. This pathway is critical for immune cell activation and explains the immunodeficiency observed in patients with mutations in either STIM1 or Orai1 (Vaeth and Feske, 2018). Therefore, the loss of SOCE in the STIM1-KO or Orai1-KO cells would be expected to decrease signaling through the calcineurin-NFAT axis in a similar fashion without necessarily affecting NFAT expression per se. So, the phenotype observed in the STIM1-KO MDA-MB-231 cells is unexpected as it leads to significant decrease in NFAT expression at the RNA and protein levels. This inhibition of NFAT expression is dependent on the loss of the STIM1 protein and not SOCE as it is not observed in Orai1-KO cells. Furthermore, NFAT downregulation in functionally significant as its reintroduction rescues the enhanced migration phenotype. These findings reveal a new function of STIM1 that is independent from its role in regulating SOCE where it maintains NFAT transcription levels at steady state and once deleted in MDA-MB-231 cells leads to a decrease in NFAT expression with the associated transcriptional remodeling that in these metastatic breast cancer cells results in enhanced migration.

STIM1 knockout in Mouse Embryonic Fibroblast (MEF) results in improved cell migration (Huang et al., 2018). Consistently, in U2OS osteosarcoma cells overexpression of a dominant-negative STIM1 mutant resulted in enhanced cell migration and larger focal adhesions (Lin et al., 2021), similar to what we observe in STIM1-KO MDA-MB-231 cells. In contrast, a STIM1 knockout U2OS osteosarcoma cell line showed decreased migration (Lopez-Guerrero et al., 2017) as is detected following STIM1 knockdown or SOCE inhibition in other cell lines (Hammad and Machaca, 2021). The reasons for these discrepancies are not clear at this point.

In conclusion, our results reveal a novel function of STIM1 that is SOCE independent and leads to NFAT transcriptional remodeling and enhanced cell migration in metastatic breast cancer cells. This needs to be considered should SOCE inhibition be considered as an anti-breast cancer therapeutic. Furthermore, the link between loss of STIM1 and NFAT transcription could be important in other cellular contexts and needs to be further investigated.

## MATERIALS & METHODS

### Cell Lines and constructs

MDA-MB-231, MCF-7, and HEK293 cells were purchased from ATCC, and were maintained in DMEM high glucose (4500 mg/L) supplemented with 10% fetal bovine serum (FBS) (AlphaFBS-HI, Alphabioregen) and 1% Penicillin/Streptomycin antibiotics (Gibco), incubated at 37 °C with 5% CO2. Retrovirus expressing NFAT1-GFP was constructed by inserting BglII/HpaI NFAT1-GFP fragment from HA-NFAT1(4-460)-GFP plasmid (Addgene #11107) into BglII/HpaI sites of pMSCV-Blasticidin plasmid (addgene #75085) and was packaged using the Phoenix-AMPHO cell line (ATCC CRL-3213). The NFAT1-GFP expressing MDA-MB-231 and STIM1-KO MDA-MB-231 cells were obtained by blasticidin selection.

### CRISPR/Cas9 Gene Editing

Single guide RNA lentiCRISPR v2 plasmids targeting STIM1, Orai1, and STIM2 were a gift from Mo Trebak (Penn State University) (Emrich et al., 2019). The sgRNA target sequences were: STIM1 g3: 5′-TGATGAGCTTATCCTCACCA-3′, Orai1 g6: 5′-GTTGCTCACCGCCTCGATGT-3′, STIM2 g1: 5′-AGATGGTGGAATTGAAGTAG-3′. Briefly, MDA-MB-231 cells were transfected with the plasmid of interest using Viromer^®^ RED following manufacturer’s instructions. After 24-48 hours of transfection, the medium was replaced by a DMEM supplemented with 10% FBS and puromycin (2μg/ml) for four passages. Clonal selection was conducted, and individual colonies validated using western blotting, sequencing (Table 1), and functional SOCE activity.

### Western blotting

Total proteins were extracted from cells using RIPA buffer (Invitrogen) supplemented with a protease inhibitor cocktail, and phosphatase inhibitor cocktail. Lysates (35-50 μg) were separated on SDS-PAGE and transferred to PVDF membrane (Bio-Rad), which were blocked with 5% milk for 1 hr and probed with primary antibody overnight at 4°C.

Primary antibodies used: STIM1 polyclonal antibody (Cell Signaling Technology #4916, 1:1000, detect extreme C-terminal end of STIM1), STIM2 (Cell Signaling Technology #4917, 1:1000), β-actin (C4) mouse monoclonal antibody (Santa Cruz #sc-47,778), Paxillin (Cell Signaling Technology # 12065), NFAT1 (Cell Signaling Technology # 5861). Both HRP-conjugated goat anti-rabbit-IgG and goat anti-mouse-IgG antibodies (1:5000) were purchased from Jackson ImmunoResearch Laboratories.

### Calcium Imaging

Cells were loaded with 1 μM Fura2-AM (Invitrogen) in Ca^2+^-containing Ringer solution at 37 □°C for 15 min, washed twice with Ca^2+^-free Ringer and imaged using a PTI imaging system mounted on an Olympus IX71. Ca^2+^ imaging data were generated from 3-4 independent experiment for each condition with an average of 60 cells per experiment. Analysis was conducted using ImageJ and Clampfit.

### Wound Healing Assay

Cells were plated in 6-well plates overnight to a confluence of about 80-90%, then the wound was created using p200 pipette tip, washed with PBS and incubated in DMEM containing 0.4% FBS. Images were taken using Inverted Fluorescence Microscope (EVOS) at different time points with the 4X objective. Waixenicin A was isolated from the soft coral *Sarcothelia edmondsoni* as described previously (Zierler et al., 2011), and its purity and identity was confirmed by ^1^H NMR (Varian Mercury Plus 300 MHz) and LCMS-ELSD (Agilent 6530 QTOF). For each experiment, 2-3 wells were examined with 4 areas of measurement in each well. Analysis was conducted using AxioVision Software (Version 4.8.2.0).

### Boydon Chamber Transwell Invasion Assay

Cell invasion was assessed using 24-well Boyden chamber (Corning). The size of the pore was 8 μm PET membrane that was coated with Matrigel. Serum-free cell suspension containing 5×10^4^ cells in 500 μl was placed in the upper chamber and 750 μl of culture medium containing 10% FBS as a chemoattractant in the lower chamber. Cells were incubated in the chambers for 13-16 hrs AT 37°C, 5% CO_2_. Cells were then fixed, permeabilized and stained with 0.5% crystal violet in methanol and visualized on a ZEISS Stemi 508 microscope. Crystal violet was dissolved in 500 μl of 10% acetic acid and absorbance recorded at 562 and 590 nm.

### Live cell tracking

Cells were seeded at low density and grown on 35 mm glass-bottom dishes (MatTek) coated with collagen I (Gibco) and cultured for 24 hrs before imaging. Time lapse live bright field imaging was performed at 37°C and 5% CO_2_ on a Carl Zeiss Laser TIRF 3 microscope using a 20× objective for 12 hrs at a rate of 1 image/3 min. Images were acquired using Zeiss Zen Blue software and ImageJ was used to manually track individual cells. Chemotaxis and Migration Tool software (Ibidi^®^) was used to plot cell tracks and compute velocity and directionality.

### Immunofluorescence Staining

Cells were fixed with 4% PFA for 15 min, followed by permeabilization with 0.1% Triton-X-100 in PBS for 10 min at room temperature. Fixed cells were blocked with 1% BSA plus 10% goat serum for 1 hr and stained with the anti-phospho-paxillin (Tyr118) antibody (Cell Signaling Technology) at 4°C overnight. Secondary Anti-Rabbit IgG (Alexa Fluor® 488 Conjugate) (Cell Signaling) was added for 1-2 hr at room temperature. Actin filaments were stained using Alexa Fluor™ 647 Phalloidin (Invitrogen). Images were captured on a Zeiss Laser TIRF 3 microscope. Images were processed and threshold using ImageJ and the FA number, size, and density was calculated accordingly.

For NFAT1 translocation experiments, cells were plated on 35-mm glass bottom dishes as described earlier and treated with and without thapsigargin, fixed, permeabilized, and stained with NFAT1 antibody (Cell Signaling Technology) and Hoechst 33342 to visualize the nuclei.

### Transcriptomics

Total RNA was extracted from MDA-MB-231, STIM1 KO, and Orai1 KO and sequenced at WCMQ Genomics Core Facility using four replicates from each cell line. Briefly, following RNA extraction, 400 ng of high integrity total RNA (RIN >5) was depleted of rRNA using the NEBNext rRNA Depletion Kit (New England BioLabs). NEXTflex Rapid Directional RNA-Seq Kit (Bio-Scientific) was used to produce strand-specific paired-end 100 bp libraries from the rRNA depleted RNA. Next, the Bioanalyzer 2100 (Agilent) on a High Sensitivity DNA chip was used to analyze the quality and quantity of the library. The libraries were then pooled and sequenced on Illumina NovaSeq 6000 in equimolar ratios. Raw sequencing data was analyzed by WCMQ Bioinformatics Core Facility. Reads were aligned and quantified with DRAGEN (Dynamic Read Analysis for GENomics) RNA Pipeline 3.6.3 (DRAGEN Host Software Version 05.021.572.3.6.3 and Bio-IT Processor Version 0×04261818) hosted on Illumina Basespace. The differential expression of genes was performed using DRAGEN Differential Expression 3.6.3 hosted on Illumina Basespace. The obtained quality–controlled reads were aligned to Homo sapiens reference genome build 38 with ensemble gene annotations using DRAGEN RNA-Seq spliced aligner. To limit the false detection rate of annotated junctions, a minimum length of 6 was used to abandon splice junctions. DESeq2 (version 1.12.4) was used to process RNA file gene count files that were produced by DRAGEN to identify the genes that were differentially expressed between 2 groups. The identified genes were further processed via calculating the log2 fold change between either STIM1-KO and MDA-MB-231 WT or Orai1-KO. The deferentially expressed genes were further processed with a significance threshold q-value of 0.05 and Log2FC over 2 up or down to identify the pathways that were differentially activated between the phenotypes.

### Proliferation Assays

Cells were seeded onto 96-well plate at a density of 5,000 cells per well. Cell proliferation was recorded at 8, 12, 24, 48, 72 hrs using the MTT (Sigma) or AlamarBlue (Invitrogen) assays according to the manufacturer’s instructions. For MTT assay, cells were processed with 5 mg/ml of MTT dissolved dimethylsulfoxide (DMSO) for 1-2 hours before each time point. Media was removed and replaced with 150μl DMSO to dissolve the formazan crystals. For AlamarBlue, the cell viability reagent was diluted down as per the recommended instructions and added 1-2 hrs before the measurement without any replacement. MTT absorbance was recorded at 570 nm against a reference wavelength at 690 nm using a microplate reader. For Alamarblue readings, fluorescence was recorded at excitation wavelength of 560 and an emission wavelength at 590 nm using CLARIOstar plus plate reader (BMG Labtech). Each group had 3 or 4 replicates along with control wells with only media to account for background absorbance or fluorescence.

### Quantitative PCR

Total RNA was extracted using RNAeasy kit (Qiagen) according to the manufacturer’s instructions. RNA was quantified using NanoDrop (ThermoFisher) before being reverse-transcribed to cDNA using High-Capacity RNA-to-cDNA Kit ((Applied Biosystems). Quantitative PCR was performed using SYBR Select Master Mix and QuantStudio™ 6 Flex Real-Time PCR System (Applied Biosystems) with the following parameter: denaturation (95°C, 2 min); 40 amplification cycles (95°C, 3 sec; 60°C, 30 sec) and melting curve analysis steps (95°C, 15 sec; 60°C, 1 min; 95°C, 15 sec). The primers used are shown in Table 2. GAPDH was used as a housekeeping gene.

**Table 2:**
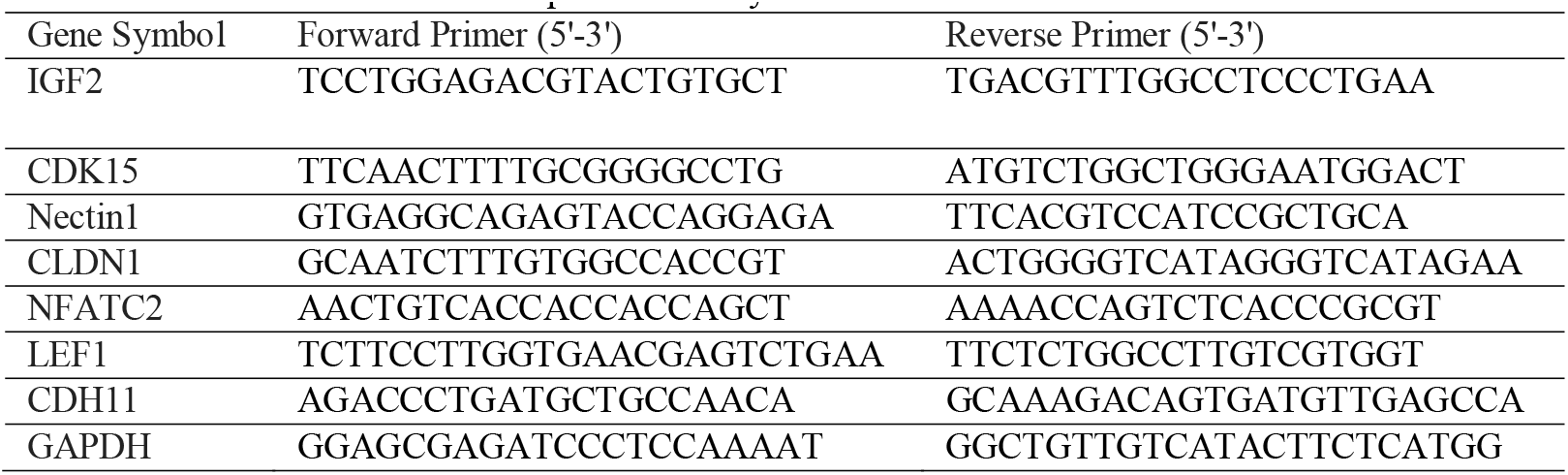
Primers Used for Gene Expression Analysis.

## Statistical Analysis

Data are reported as Mean ± SEM with statistical analyses performed using Prism (GraphPad) with the statistical test indicated. For all tests, *p<0.05; **p<0.01; ***p<0.001; ns: not significant.

## ACKNOWLEDGEMENTS

We thank the Microscopy Core at WCMQ for helping in experiments. The Core is supported by the ‘Biomedical Research Program at Weill Cornell Medical College in Qatar’, a program funded by Qatar Foundation. The statements made herein are solely the responsibility of the authors.

## Data Availability

All data generated are included in the manuscript and supporting data.

